# Facial and body colouration is linked to social rank in the African cichlid *Astatotilapia burtoni*

**DOI:** 10.1101/2024.08.21.608989

**Authors:** M. Peroš, A. Chang, A. Martashvili, S.G. Alvarado

**Affiliations:** City University of New York Neuroscience Collaborative Subprogram, Graduate Center of City University of New York, New York, NY 10016; Subprogram in Ecology, Evolution, and Behavior, Department of Biology, Graduate Center of City University of New York, New York, NY 10016; Cognitive Neuroscience Program, Graduate Center of City University of New York, New York, NY 10016; Department of Biology, Queens College City University of New York, NY 11367

## Abstract

Animal colouration is fundamentally important for social communication within conspecifics to advertising threat to competitors or fitness to possible mates. Social status and animal colouration are covarying traits that are plastic in response to dynamic environments. In the African cichlid, *Astatotilapia burtoni*, body colouration and behaviour have been extensively reported to vary with social rank. However, the nature of the interaction between these two traits is poorly understood. We hypothesise that pigmentation patterns could be linked to the behavioural repertoires underlying social status and can be resolved to regions on the cichlid body plan. To test this hypothesis, we generated Territorial (T) and Non-territorial (NT) males and employed computer vision tools to quantify and visualise patterns/colour enrichment associated with stereotyped T/NT male behaviour. We report colour-behaviour interactions localised in specific areas of the body and face for two colour morphs illustrating a more nuanced view of social behaviour and pigmentation. Since behavioural and morphological variation are key drivers of selection in the East African Great Rift Lakes, we surmise our data may be translatable to other cichlid lineages and underline the importance of trait covariance in sexual selection and male competition.

## Introduction

Animal colouration is a fundamentally important trait across the animal kingdom. Diversity and plasticity through pigmentation serve various functions, from thermoregulation to social signalling and cryptic displays. For example, Balkan moor frogs [1] and stony-creek frogs [2] undergo dramatic changes in body colouration during mating, while bearded dragons modulate dorsal pigmentation for crypsis and thermoregulation in the wild [3]. The development and plasticity of animal pigmentation are critical to the diversification and adaptive speciation of various animal clades. For example, industrial melanism in peppered moths is a classic example of how natural variation and selection can shape population dynamics [4]. Across taxa, the evolution of unique colours and patterns is linked to ecology and antipredator defences related to background adaptation [5] and intermediary phenotypes to evolve into warning colouration [6]. The biological basis for colouration is often resolved to cellular and molecular substrates, which underline the importance of developmental genetics in studying animal pigmentation. However, animal colouration is also plastic in response to various social, physical, and chemical stimuli comprising the ambient environment [7].

The link between animal colouration and behaviour has been studied for nearly a century and illustrated for roles in sexual selection and social communication. The evolution of nuptial colouration in birds and fishes has been shown to drive mate selection and competition [8]. However, these roles expand beyond sexual selection and can result in additional pleiotropic effects. For example, agouti signalling is fundamentally essential for melanin deposition in mammals but has also been shown to modulate social [9] and feeding behaviours [10]. Similarly, stimulation of melanocortin signalling in cichlid fish can regulate pigment cells’ function and shape aggressive behaviour in males [11]. The link between animal behaviour and pigmentation is vital concerning phenotypic plasticity in heterogeneous environments. For example, seasonal variation can shape visual ecology, territory establishment, and subsequent mating opportunities critical for survival[12]. However, most of the existing literature has focused on resolving animal pigmentation to genetic variation instead of developing an understanding of phenotypic plasticity in animal colouration and behaviour.

Within cichlid lineages, morphological variation and behaviour have long been considered key factors driving their adaptive speciation [7,13]. Sensory drive suggests that interactions between sensory systems and the ambient environment can shape evolution on predictable trajectories and are salient for animal colouration and perception [14]. For example, variation in the nuptial colouration of cichlid *Pundamilia* lineages has been linked to selection acting on the *lws* opsin along all but the steepest environmental gradients within an island in Lake Victoria. In contrast, eutrophication and increases in the turbidity of Lake Victoria have been suggested to lead to species collapses where intraspecific visual signalling can be occluded [15,16]. Despite the importance of animal pigmentation in diverse processes, the intersectional roles of behaviour and animal pigmentation still need to be better understood, suggesting a nuanced study of behaviour and animal pigmentation is merited. While developmental genetics has identified the molecular bases of colouration/behaviour within populations [17–19], it has not been as helpful in understanding the plasticity of these inherently dynamic model traits.

Here, we adopt the African cichlid model system, *Astatotilapia burtoni*, for its robust social system [20] and morphological variation [11,21–24]. *A. burtoni* has a robust social hierarchy and the ability to change colour, which allows proper investigation of how behaviour and colour might be linked. In nature, males become territorial (T), displaying brighter colouration [25], black eyebars [26,27], and egg spots on the anal fin, whereas non-territorial (NT) males are muted in colour and do not hold territory. These social ranks and accompanying patterns are reversible via social descent and ascent paradigms[28,29] and present in both blue and yellow male colour morphs, each with their behavioural repertoire [21,30]. For example, yellow individuals preferentially interact with individuals of the opposite colour and are better able to establish and defend territory than blue males [23]. Additionally, yellow morphs are more aggressive and win more fights against blue morphs in direct territorial competitions [11].

In this study, we manipulated social status and male colouration to investigate the colours associated with social behaviour and interindividual signalling. Males were reared in a dyad paradigm to generate T and NT males in the presence of three females on either blue/yellow substrates with downwelling light of the same colour. Dyads were filmed, photographed, and manually scored. On photographs of the males, we applied a k-means reduction of colouration in fish to quantify colour pattern areas and correlative analyses to evaluate if body colouration is associated with behavioural variation. Colours linked to specific behaviours were then mapped onto cichlid body plans to identify underlying colour distribution patterns unique to social rank. Cichlids’ conspicuous and cryptic colours are important for sexual selection and male-specific competition and are relevant for understanding interindividual signalling. Further, the blue-yellow colour morph dualism, the bright-muted colours of T/NT males, and secondary patterns such as pelvic fin stripes, humeral blotch, and egg spots are shared across various lineages of cichlids in the East African Great Rift Lakes [31,32]. These data will contribute to understanding the interactions between plastic traits in a model family that are important for the largest adaptive speciation seen in vertebrates.

## Methods

### Animal husbandry and dyad tank setup

Stockfish were reared in communal settings under laboratory conditions mimicking their natural environment: pH 7.8 ± 8.2, 29°C, and a 12-hour light/dark cycle with full-spectrum illumination[33]. Water was regulated using an automated sensor system (Neptune Systems, Apex) to dispense salt (Seachem, #0279) and buffer (Seachem, #0287). Animals were housed on a brown gravel substrate (Pure Water Pebbles #30035) with halved terra-cotta pots to allow males to establish and maintain their territories [34] and fed daily at 9:00 AM with flakes (Tetra AQ-77007). Tank gravel colour has previously been sufficient to reversibly induce distinct colour phenotypes in *A. burtoni [35]*. To determine the effect of environmental colour on social behaviour, dyad paradigm tanks were supplied with yellow (Pure Water Pebbles, #70021) or blue gravel (Pure Water Pebbles, #70111) with matching downwelling light from custom-made LED lighting. A dyad of two socially naive, size- and age-matched males and three females were selected and placed into either yellow (n=18) or blue (n=28) tanks for four weeks to establish a social hierarchy, generating T and NT phenotypes.

### Behavioural analysis, photography, and animal sacrifice

Tanks were recorded using video cameras (GoPro Hero8 Black) mounted on aluminium tripods (AmazonBasics WT3540). Tripods were positioned so camera lenses were ∼15 cm away from the fish tank, and height was adjusted until the whole tank was visible (∼105cm). Video recordings were initiated at noon under “Standard” settings (1080p, 30FPS, linear) for 60 minutes. Videos were manually scored with BORIS event-logging software [36] utilising a constructed ethogram described in Supplemental Table 1 with behaviours described in previous literature[25,33]. Scoring began after a 15-minute habituation period. Then, males were collected and pat dry with a paper towel for photography in a lightbox (Puluz PU5032B) next to an X-Rite colour standard (X-Rite MSCCMN-RET). Animals were weighed using a digital scale (±0.001g) and lengths recorded for mouth-to-caudal trunk before being sacrificed via rapid cervical transection (IACUC Protocol #198). Chemical/thermal anaesthetics were avoided here to mitigate any impact on melanophore/chromatophore expression under nervous control [27]. Following photography and sacrifice, gonads were dissected and weighed to calculate the gonadosomatic index (GSI) as a function of reproductive investment.

### Image analysis

Photographs were colour-corrected (Fig. 1A) using the colorChecker function from the R package ‘patternize’ [37]. White light reflected on eyes resulted in overexposure in all images. At the same time, fin positions varied picture-to-picture and were not equally splayed to observe patterning. Following correction, overexposed pixels, fins, and eyes were masked to a black background using FIJI (ver. 2.15.1) to avoid type I errors in colour analysis (see Fig. 1B, left side) [38]. To segment colour patterns between colour morphs and quantify the relative area covered by specific colours, the ‘recolorize’ package was used to carry out a k-means clustering of all colours in all images to a palette of 30 colours for each colour-rearing group (Fig. 1C) for remapping on each image [39]. To reduce colour variation, the photos were blurred using the ‘blurImage’ function in recolorize. The 30-colour palette was chosen as it represented the most straightforward and accurate colours from the original images (see Fig. 1B). Due to discrete differences in blue/yellow male colour morphs (Fig. 1A), two different colour palettes were used for yellow and blue male morphs (Fig. 1C). This analysis guided false colouring of colours that exceeded a 5% threshold for total area (see Figs. 2 and 4). colours that significantly correlated with social rank were then remapped using false colours to examine patterns of colour seen across individuals as heatmaps via ‘patternize’ for the whole body and faces.

**Figure 1:**
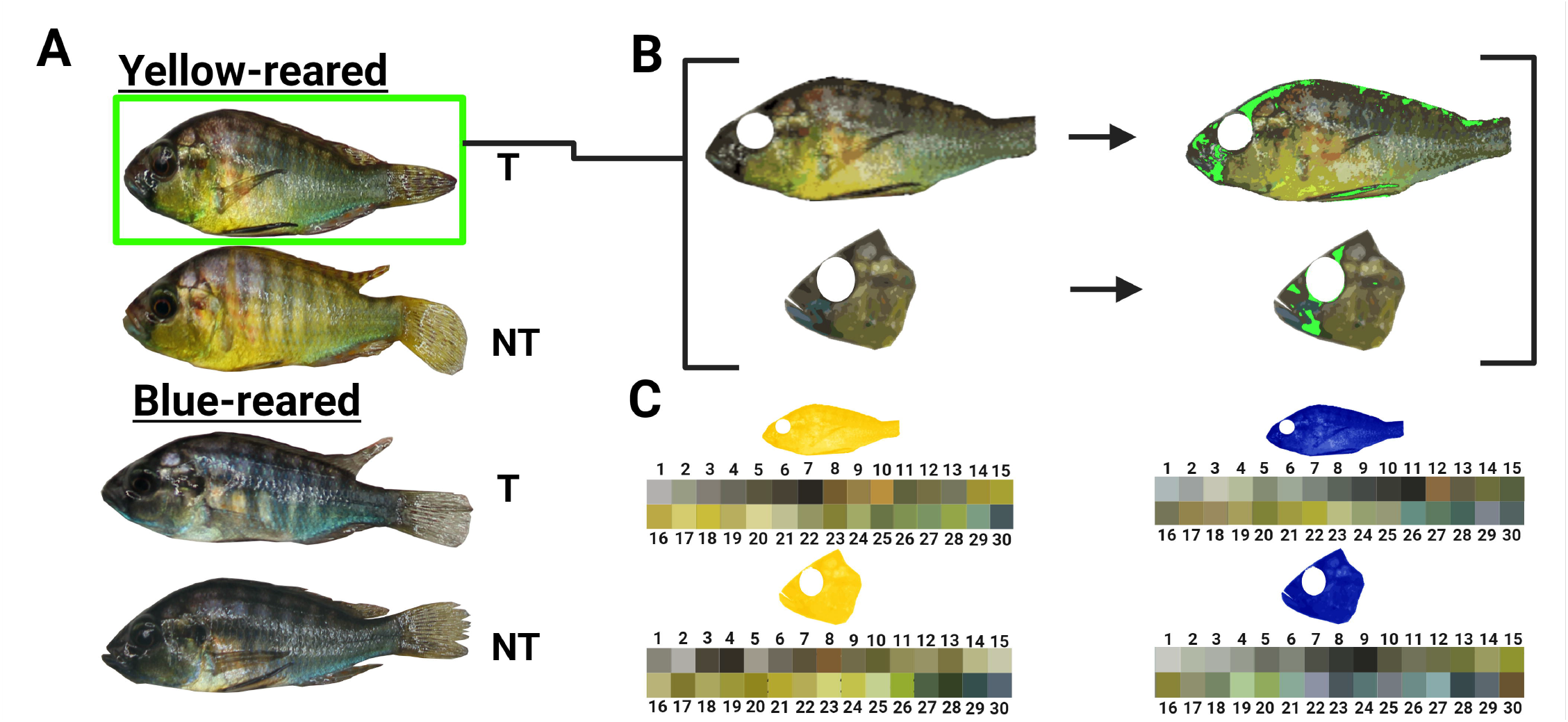
Computer vision workflow used to analyse body colour patterns in A. burtoni territorial and non-territorial males (A) Example images of yellow- and blue-reared territorial (T) and non-territorial (NT) males used in this experiment (B) Images were masked to isolate either the whole body or face (left side) for later colour analysis using false colour highlights (right side). (C) Colour palettes generated using recolorize for rearing environment (left-right) and body segment (top-bottom).

**Figure 2:**
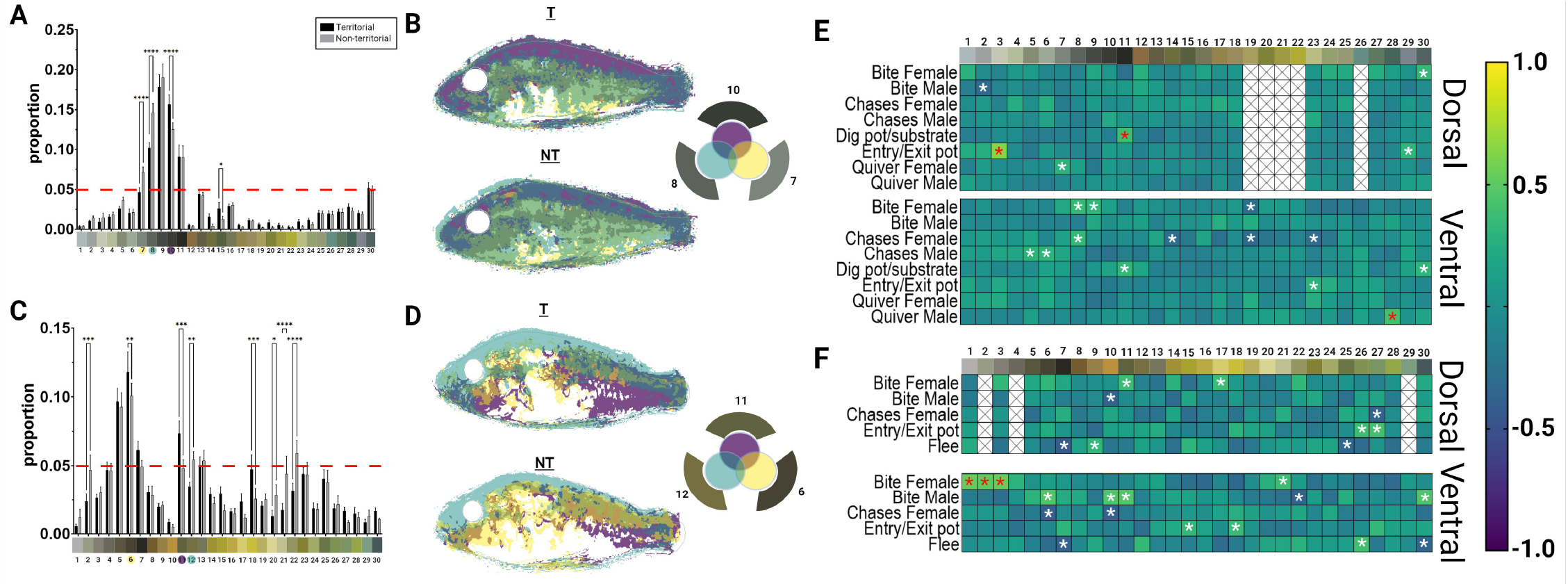
Quantification and patterning of body colour and correlation to behaviour in blue- and yellow-males. Quantification of colour variation in blue-(A) and yellow-reared (C) males. Mean plotted with SEM as error bars, N_blue_=28, N_yellow_=18, One-way ANOVA, Tukey’s post hoc. (B) Top three colours in (A) layered as a composite overlay used to falsely colour the cichlid body plan separately with rank in blue-reared males. (D) Top three colours in (C) layered as a composite overlap used to falsely colour the cichlid body plan separately with rank in yellow males. Outer colours on Venn diagram inset represent significant colours falsely coloured on the inner Venn diagram. ** p <0.07, ***p<0.007,****p<0.0007. (E,F) Correlations table between behaviour and dorsal/ ventral colour. Blue-reared (E) and yellow-reared (F) males. Each cell represents a Pearson’s correlation coefficient (r), with yellow and blue colour coding positive and negative r values respectively. White, crossed boxes represent absent values for colours on ventral/ dorsal halves.

### Statistical analysis

Score logs exported from BORIS software were analysed in RStudio-2023.12.1-402, using base R and packages such as ‘stats’, ‘dplyr’, ‘ggplot2’, ‘tidyr’, and ‘patchwork’ [40]. Relative frequencies were calculated, normalising behaviours to total behaviours within each subject. Two-way ANOVA was conducted using GraphPad Prism v. 10.2.2. Paired multiple comparisons tests were performed on each behaviour category using the Tukey multiple comparisons test. GraphPad Prism was used for colour analysis to quantify the variation of morph-specific 30-colour bins. A two-way ANOVA was conducted, followed by Tukey multiple comparisons test. GraphPad Prism was also used to generate multiple correlation tables for colour and behaviour on the dorsal and ventral halves of the cichlid body plan. Significant correlations were individually plotted against relevant behaviours for social rank for body and face in each colour morph, where we reported Pearson’s *r, r*^*2*^, and 95% confidence intervals on the data.

## Results

### Dorsal and ventral body colour patterns correlate with behaviour and colour morph

To examine the interaction between body colouration and behavior, we simplified the number of colours represented by pixels within each fish image and remapped those colours back onto the fish. This generated images that recoloured original images (Fig. 1A) into 30-colour palettes for each rearing environment (Fig. 1B and C). Pixel areas of each colour were quantified within ‘recolorize’ per individual and normalised as within-individual proportions by dividing colour pixel area by the sum of all colour pixels (Fig. 2A and C). Our ANOVA showed that among blue-reared males, colour was responsible for 68.28% of the total variance (Fig. 2A, F_colour_(29, 1500)=116.1, p <0.0001; F_rank_(1, 1500) = 2.21 ×10^−11^, p >0.99; F_interaction_(29,1500) = 2.19, p<0.001). Three colours were significantly different between ranks (Tukey post hoc, p<0.0001). Among yellow-reared males, colour was also responsible for most of the variation (Fig. 2C, F_colour_(29,1020)=29.16, p <0.0001; F_rank_(1, 1020) = 1.82 ×10^−11^, p >0.99; F_interaction_(29,1020) = 2.16, p<0.001). Four colours were significantly different between ranks (Tukey post hoc, p_6_<0.01, p_11_<0.001, p_12_<0.01, and p_22_<0.0001). Three of these colours were then mapped using false colours representing variation in both dorsal and ventral areas of the body (Fig. 2B and D). In the case of yellow-reared males, the colours with the greatest magnitude of Pearson’s *r* were selected (Fig. 2F). In addition to our focus on rank (Fig. 1A-D) we decided to carry out a correlation analysis across all animals involved in the study categorized by their rearing colour (Blue: Fig1E or yellow: Fig1F). Since *A. burtoni* are typically found in shallow areas in Lake Tanganyika, they are likely to develop cryptic colouration on the dorsal portion of their bodies and display conspicuous, socially relevant colours on the ventral portion [41]. Each image’s pixel area proportions were separated into dorsal and ventral portions to account for the biological significance of midline divisions. To test whether colour pattern areas correlate with behaviour, Pearson’s multiple correlation matrices were generated for each division, with significant correlations adjusted for multiple comparisons (Fig. 2E and F).

For each rearing colour, the top three significant correlations were selected based on Pearson’s *r* magnitude (Figure 2E and F, red asterisk). Among blue-reared males, three different behaviours (*Entry/exit pot, Dig pot/substrate*, and *Quiver male*) correlated with colours while three colours correlated with *Bite female* in yellow-reared males. For each correlation, a rank-independent colour heat map visualises pattern similarities within rearing environment while rank-dependent scatterplots were used to determine rank-specific contributions to the compound Pearson’s *r* in Figure 2 (Figure 3). In blue-reared males (Fig. 3A), *Entry/exit pot* correlated with colour 3, which is ventrally expressed with areas of relatively low pattern agreement largely driven by a positive correlation in T males (*R*^*2*^_T_ = 0.307, *R*^*2*^_NT_ =0.073). *Dig pot/substrate* correlates with colour 11, dorsally located with an area of strong pattern agreement, but did not correlate with either rank (*R*^*2*^_T_ = 0.089, *R*^*2*^_NT_ =0.018). *Quiver male* correlated to colour 28, which shows ventral pattern agreement towards the caudal trunk, was positively correlated in T males (*R*^*2*^_T_ = 0.361, *R*_NT_ ^*2*^ =0.049). In yellow-reared males (Fig. 3B), *Bite female* correlated with three similar colours that exhibit a diagonal dorsal-to-ventral pattern. colours 1 and 2 both positively correlate in NT males (**colour 1** *R*^*2*^_T_ = 0.0811, *R*^*2*^_NT_ =0.484; **colour 2** *R*^*2*^_T_ = 0.0127, *R*^*2*^_NT_ =0.453) while colour 3 positively correlates with both T and NT behaviour (*R*^*2*^_T_ = 0.202, *R*^*2*^_NT_ =0.290).

**Figure 3:**
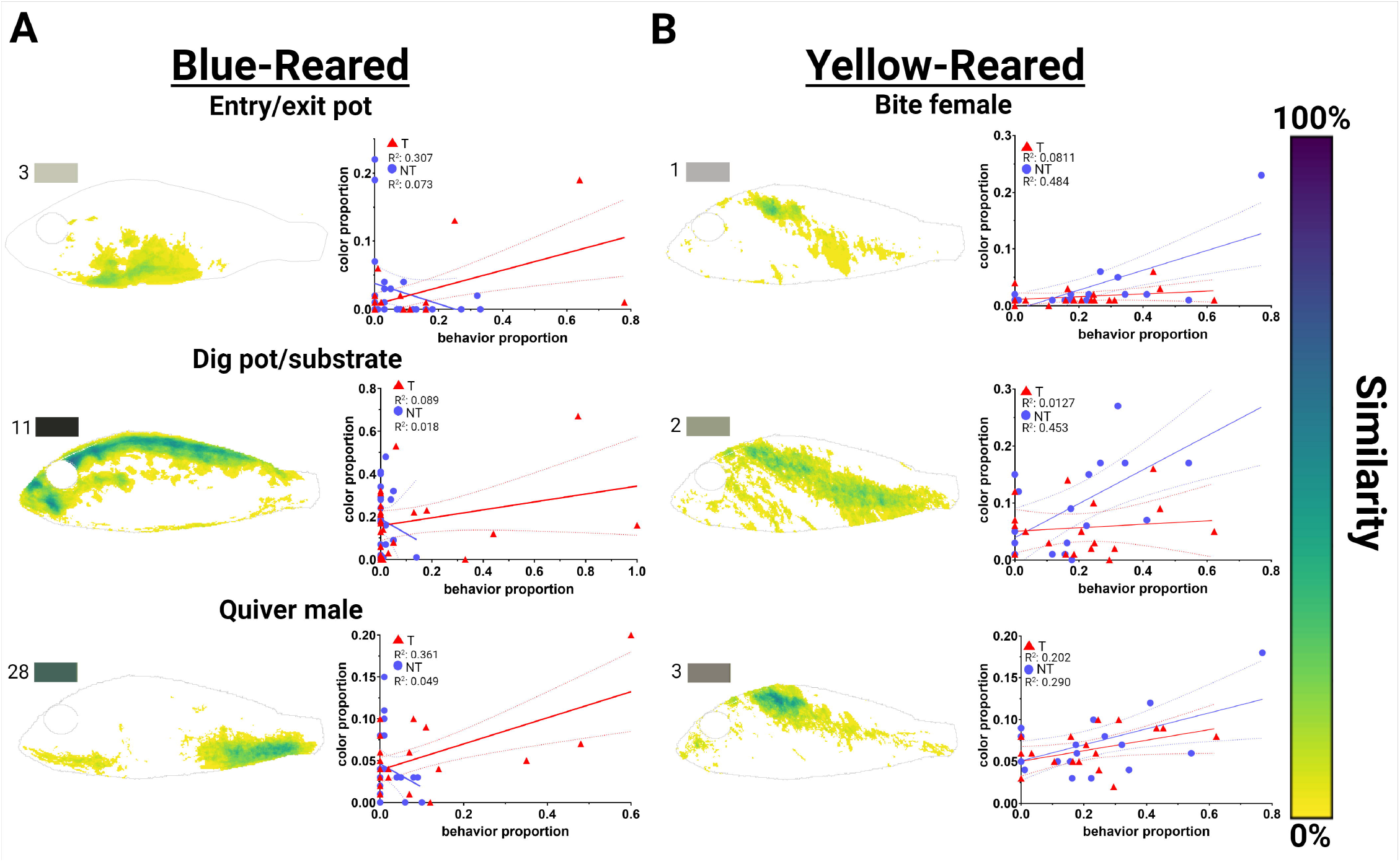
Colour patterns and rank-dependent scatterplots of top three behaviour-colour correlation pairs. Colour area agreement and rank-dependent correlations of blue-(A) and yellow-reared (B) males. Heatmaps (left) indicate low (yellow) or high (dark blue) levels of pattern overlap among all blue-reared males for the indicated colour. Scatterplots (right) showing individual values for colour and behaviour proportions in territorial (T, red triangle) and non-territorial (NT, blue circle) males. Solid and dotted lines represent best flt line and 95% confidence intervals respectively.

**Figure 4:**
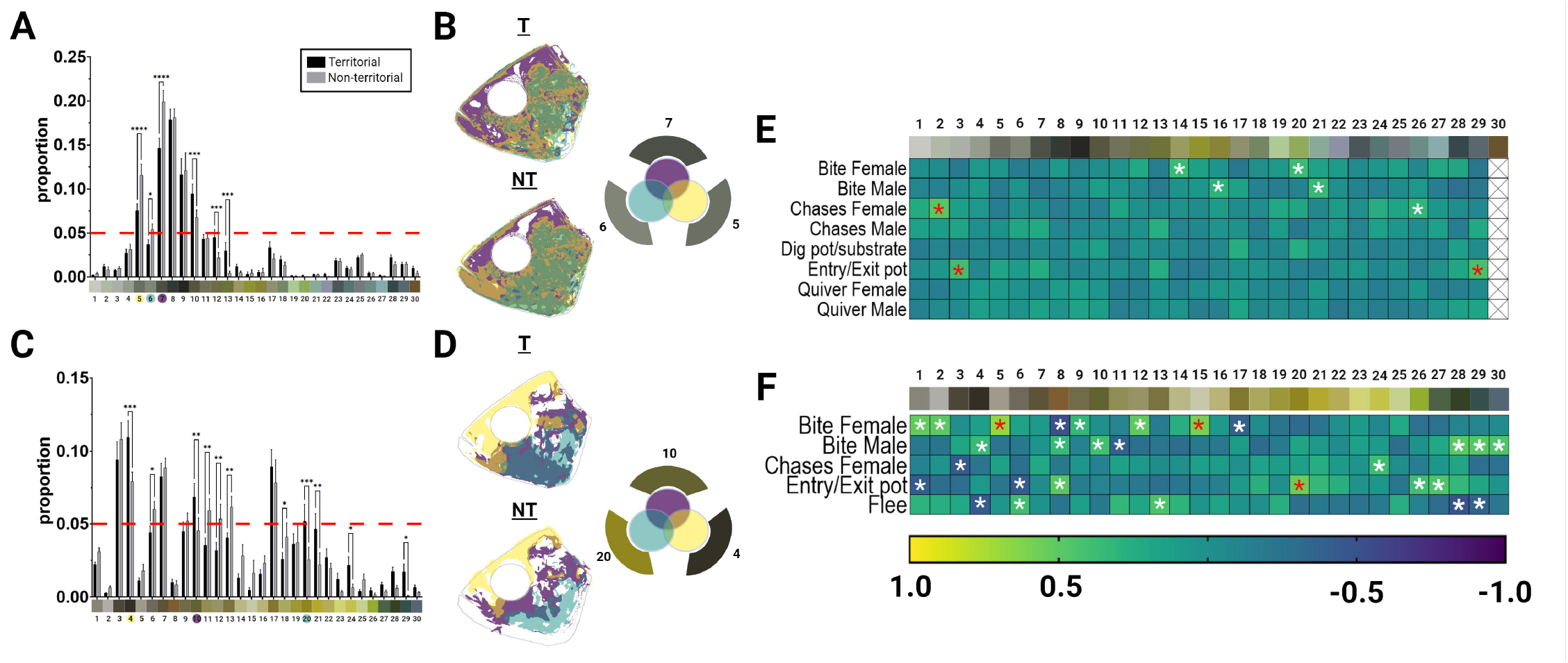
Quantification and patterning of face colour and correlation to behaviour in blue- and yellow-males. Quantification of colour variation in blue-(A) and yellow-reared (C) males. Mean plotted with SEM as error bars, N_blue_=28, N_yellow_=18, One-way AN OVA, Tukey’s post hoc. (B) Top three colours in (A) layered as a composite overlay used to falsely colour the cichlid face separately with rank in blue-reared males. (D) Top three colours in (C) layered as a composite overlap used to falsely colour the cichlid face separately with rank in yellow males. Outer colours on Venn diagram inset represent significant colours falsely coloured on the inner Venn diagram.** p <0.01, ***p<0.007, ****p<0.0001. (E,F) Correlations table between behaviour and colour. Bluereared (E) and yellow-reared (F) males. Each cell represents a Pearson’s correlation coefficient (r), with yellow and blue colour coding positive and negative r values respectively. White, crossed boxes represent absent values for colours on the face.

### Proportions of facial colour correlate with social behaviour

Previous literature suggests facial features are a key visual focal point for fish [42]. To examine which facial colour patterns correlate with behavior, we isolated the anterior portion of the fish images that contained the face and quantified patterns of colour that were associated with social behavior. In blue-reared males, colour is responsible for 67.95% of the variance in the data (Fig. 4A, F_colour_(29, 1500)=116.05, p <0.0001; F_rank_(1, 1500) = 7.44 ×10^−5^, p >0.99; F_interaction_(29,1500) = 3.01, p<0.0001). Of colours that differed between ranks, four colours with proportions above 5% are significantly different between ranks (Tukey post hoc, p_5_<0.0001, p_6_<0.05, p_7_<0.0001, and p_10_<0.001). Colours plotted in the heatmap (Fig. 4B) had the three greatest Pearson’s *r* magnitudes, excluding colour 10. Among yellow-reared males, colour is responsible for 49.31% of the variance in the data (Fig. 4C, F_colour_(29, 1020)=36.70, p <0.0001; F_rank_(1, 1020) = 9.77 ×10^−9^, p >0.99; F_interaction_(29,1020) = 2.54, p<0.0001). Of colours that differed between ranks, seven colours with proportions above 5% are significantly different between ranks (Tukey post hoc, p<0.05). colours plotted in the heatmap (Fig. 4D) had the three greatest Pearson’s *r* magnitudes.

For each rearing colour, three significant correlations were selected based on Pearson’s *r* magnitude (Figure 4E and F, red asterisk). Two different behaviours for blue (*Chase female* and *Entry/exit pot*) and yellow (*Entry/exit pot* and *Bite female*) correlate with colours. For each correlation, a rank-independent colour heat map visualises pattern similarities within rearing environment while rank-dependent scatterplots were used to determine rank-specific contributions to the compound Pearson’s *r* in Figure 4 (Figure 5). In blue-reared males (Fig. 5A), *Chase female* correlated with colour 2, expressed with areas of relatively low pattern agreement and little rank-specific correlation (*R*^*2*^_T_= 0.045, *R*^*2*^_NT_=0.014). *Entry/exit pot* correlates with colours 3 and 29 (**colour 3** *R*^*2*^_T_= 0.049, *R*^*2*^_NT_=0.262; **colour 29** R^2^_T_= 0.043, R^2^_NT_<0.001). In yellow-reared males (Fig. 5B), *Entry/exit pot* correlates with colour 3, driven by T male correlation (*R*^*2*^_T_= 0.558, *R*^*2*^_NT_=0.002). colours 1 and 2 both have strong positive correlations in NT males (**colour 1** *R*^*2*^_T_= 0.222, *R*^*2*^_NT_=0.652; **colour 2** *R*^*2*^_T_= 0.055, *R*^*2*^_NT_=0.558).

**Figure 5:**
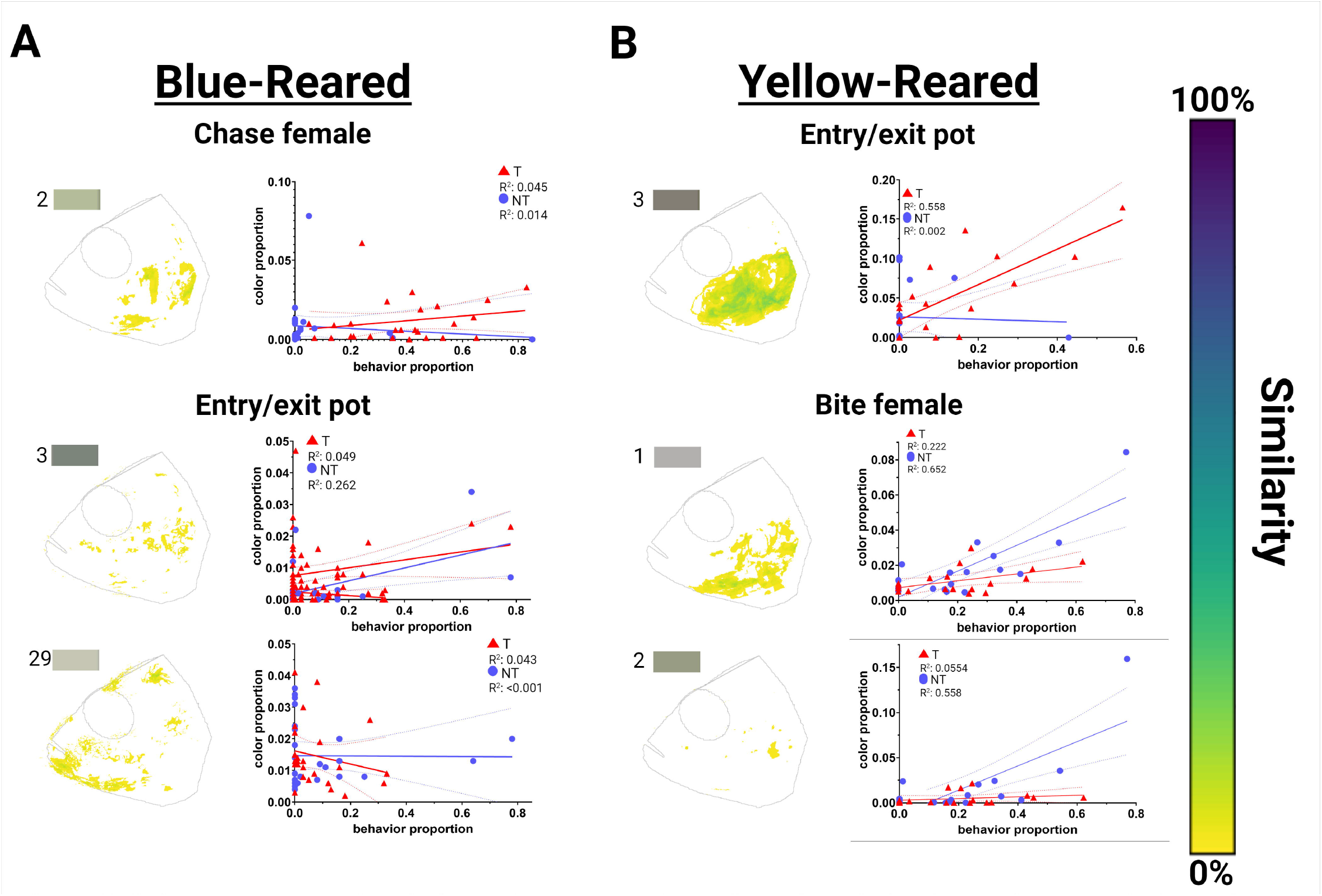
Colour patterns and rank-dependent scatterplots of top three behaviour-colour correlation pairs specific to the cichlid face. Colour area agreement and rank-dependent correlations of blue-(A) and yellow-reared (B) males. Heatmaps (left) indicate low (yellow) or high (dark blue) levels of pattern overlap among all blue-reared males for the indicated colour. Scatterplots (right) showing individual values for colour and behaviour proportions in territorial (T, red triangle) and non-territorial (NT, blue circle) males. Solid and dotted lines represent best fit line and 95% confidence intervals respectively.

## Discussion

African cichlids in the East African Rift Lakes have long been recognized as textbook examples of sexual selection, with morphological diversity being a driving factor steering their adaptive speciation[13]. Within this family, *Astatotilapia burtoni* is emerging as a powerful model for studying sociobiology[20] and phenotypic plasticity[7]. In this study, we generated socially dominant territorial (T) and subordinate non-territorial males (NT) of two known colour morphs. We tested the hypothesis that quantitative variation in colouration is linked to modalities of social rank and behaviour. First, we show that different colour morphs have unique palettes of colour variation across social rank. Secondly, we show that these colours can vary with behaviour independent of rank. Lastly, we localise this variation across the cichlid body plan and face. Together, these data provide a descriptive overview of two colour morphs present within this species that may reveal how and where visual signalling occurs within the social hierarchies of African cichlids.

*A. burtoni* has a robust social system in which aggressive, conspicuously coloured T males hold territory and court females, and subordinated, cryptic NT males have none of these opportunities. Males of either rank can express blue or yellow body colours in the wild and the laboratory as a function of social community [21] and/or visual ecology [22,35]. However, it is unclear if body colouration is expressed as a function of social life history or other variables. In previous work[21–23], the age of individual males was not controlled, which may shape the results of social competition across male size differentials [43]. Here, body colouration was controlled with age-matched males that were never before dominant and paired within a dyad paradigm in a blue or yellow visual environment. Thus, animal colouration was solely induced by the ambient light in the rearing environment where social life history and age were controlled. It is important to note that this approach also removes contrasting colour pairs between T/NT dyad males from all analyses [23]. We suspect this manipulation through the visual environment is more analogous to seasonal shifts in visual ecology seen in Lake Tanganyika, where colour morphs are localised regionally along the shoreline (unpublished data).

Various physiological cues are capable of shaping variation in colouration and rank. For example, the expression of endothelin receptors in yellow pigment cells has been shown to modulate the dispersal of pigments through epigenetic substrates[35] in blue and yellow morphs. Additionally, both yellow and blue male colour morphs have unique hormonal profiles when controlled for size (but not age or visual environment). Higher levels of plasma 11-ketotestosterone were measured in yellow T males compared to blue T males, while blue NT males had higher plasma cortisol [21] or differences in their melanocortin system (Dijkstra 2017). Similarly, androgen signalling may also play an upstream role in how colouration is expressed as a function of social behaviour. In androgen receptor knockouts, male fish show muted colouration across their body, suggesting a deficiency in developmental pigmentation[44] analogous to albino tyrosinase knockouts within *A. burtoni*[45].

Ultimately, the colours and patterns described in this study can be resolved down to cellular substrates, chromatophores, which function by rapidly aggregating/dispersing pigment granules in response to both neural and physiological cues. For example, phenotypic plasticity in the body coloration of *Melanochromis auratus* has been attributed to the innervation of dendritic processes around melanophores of their black morphs when compared to yellow morphs[46]. However, this study could not report on the neural and physiological control of chromatophore function (or its associated patterns). Here, morphological variation in these colours was reported as the presence or absence of colours and patterns in the dermal chromatophore unit. When handled for photographs, animals were subject to brightening through largely dispersed chromatophores. This colouration and its change occurs over weeks relying on changes in the density, function, and overlapping processes of chromatophores of different pigments[7]. Furthermore, while we may identify pixel variation for dark or light yellow, it should be assumed that this colouration is tied to the interactions of yellow xanthophores, melanophores, and iridophores in specific locations on the cichlid body plan for an individual pixel.

Across the cichlid body plan quantitative variation in coloration was observed in both colour morphs and with rank. False colouring highlighted overlapping localization of different colours were most notable in blue males. Specifically, blue males had two overlapping dark tones (Colour 8 + 10, Figure 2B) covering the dorsal side compared to yellow males who had mostly yellow tones on their dorsal side and throughout their body (Figure 2D). This dorsal darkening was also identified to be correlated with digging within the pot territory in blue males, albeit with low R^2^ values (Figure 3A). Blue males have been suggested to more sexually attractive in female mate choice experiments[47] and we suspect that female preference for blue males may also be tied physiologically induced changes in opsin expression observed in the eyes of female *A. burtoni* and an increased sensitivity to male colouration [48] (See Colour palettes generated in Figure 1C). Specifically, the increased expression of the short wavelength opsins (sws1/2a/2b) in ovulated females are sensitive to light within the violet-blue range [48] highlighting blue males. Within blue males, a blue-green patch (Colour 28, Figure 3A) was identified in the ventral posterior end revealing a correlation with male quivering mainly within T males (R^2T^= 0.361 vs R^2NT^=0.05). Between males, quivering has been interpreted as a threat display between males and often involves aligning bodies where the posterior third of the quivering males overlaps with the visual field of the observing male. Since the rh2b opsin mRNA has been reported to be enriched in NT males it may allow for a more vivid perception of green across this patch in T males between signaller (T males) and receiver (NT males). However additional experiments will need to be carried out to consolidate this link. In general, our data suggest that brighter contrasting colours are present in T compared to NT males. We suspect it may be due to sexual selection and female mate preference. This is analogous to how androgen receptor knockouts, who display developmentally muted coloration, are less sought after by females (Howard 2023).

Prominent facial features exclusive to T males include a dark bar pattern under the eye, which is suppressed in NT males and can be turned off rapidly [26]. For example, cutting the Vth cranial nerve or the administration of norepinephrine can discretely attenuate the black eyebar through the rapid aggregation of underlying melanophores. In our data, colours correlated with social behaviour were localised to patterns enriched in the face of *A. burtoni* (Figures 2 and 4). In the face, this was seen as darker tones on the dorsal anterior face in T compared to NT males that spread further towards the ventral lip and jaw (Figure 4BD). Across yellow males, grey tones were seen to be correlated with both digging and chasing females and were largely localised anterior to the operculum (Figure 5B) and were most correlated among T males compared to NT counterparts. We speculate that this pattern may be tied to threat behaviours such as frontal displays where competing males face each other with open jaws typically leading to demarcated boundaries between territories[25,33]. This is supported by behavioural variation we have observed in socially naive yellow (not blue) males(unpublished data) and how these patterns are enriched in the “cheek” via false coloured correlations (Figure 5A).We suspect that this colour variation in the face supports other evidence in cichlid fish where attention on the face may signal intraspecific social signalling[42,49]. For example, other Tanganyikan cichlids, such as *Neolamprologous brichardii*, focus more on the faces of conspecifics when presented with images bisecting the body plan of conspecifics/heterospecifics[42]. Similarly, *Neolampologous pulcher* focuses more on familiar conspecifics than strangers[49].

Our data contextualise how the phenotypic plasticity of male stereotyped coloration may send information to conspecifics based on rank and/or sex. To date, studies on morphological variation in pigmentation focus wholly on developmental variation in pigmentation as a function of genetic variation or qualitatively categorise pigmentation as the absence/presence of patterns or colours. Furthermore, they rarely address developmental plasticity in adults. Here, computer vision workflows afforded us a more nuanced and quantitative study of pigmentation across two colour morphs in a phenotypically plastic model system. Since markings such as anal fin spots, eye bars, pectoral fin striping, red humeral blotch, and high contrast striping are found across various cichlid lineages, we anticipate that the data presented here may be translatable to other cichlid and teleost models. These data can also inform how exogenously added markings either via latex injections or tattoos can allow to testable hypotheses on signal-receiver dynamics and social communication. Since these findings sit at the crux of behavior and morphology, two known drivers of selection in the East African Great Rift Lakes, they may allow a deeper understanding of the covariation of traits contributing to evolutionary outcomes.

## Supporting information

Supplemental Table 1. Description of all behaviours scored with mean frequency and SEM.

## Acknowledgments

We would like to thank Alvarado Lab members Joseph Lawrence, Yael Sassoon, and Elana Anavian for collecting animals used for photography in this study. Thanks to Yael Mushell, Robert Ilyasov, Daniel Iskhakov, and Sofia Taherkhani for carrying out behavioural scoring in this study. We also would like to thank Yefim Radomyselskiy for developing the overhead lighting system to change the colour of the rearing environments and Dr. Maral Tajerian for her feedback and comments on this manuscript and project.

## References

1. Ries C, Spaethe J, Sztatecsny M, Strondl C, Hödl W. 2008 Turning blue and ultraviolet: sexLJspecific colour change during the mating season in the Balkan moor frog. J. ZooDl. 276, 229–236. (doi:10.1111/j.1469-7998.2008.00456.x)

2. Kindermann C, Narayan EJ, Hero J-M. 2014 The Neuro-Hormonal Control of Rapid Dynamic Skin Colour Change in an Amphibian during Amplexus. Plos One 9, e114120. (doi:10.1371/journal.pone.0114120)

3. Smith KR, Cadena V, Endler JA, Kearney MR, Porter WP, Stuart-Fox D. 2016 Color Change for Thermoregulation versus Camouflage in Free-Ranging Lizards. Am. Nat. 188, 668–678. (doi:10.1086/688765)

4. Kettlewell HBD. 1955 Selection experiments on industrial melanism in the Lepidoptera. Heredity 9, 323–342. (doi:10.1038/hdy.1955.36)

5. Hoekstra HE. 2006 Genetics, development and evolution of adaptive pigmentation in vertebrates. Heredity 97, 222–234. (doi:10.1038/sj.hdy.6800861)

6. Loeffler-Henry K, Kang C, Sherratt TN. 2023 Evolutionary transitions from camouflage to aposematism: Hidden signals play a pivotal role. Science 379, 1136–1140. (doi:10.1126/science.ade5156)

7. Alvarado SG. 2020 Molecular Plasticity in Animal Pigmentation: Emerging Processes Underlying Color Changes. Integr Comp Biol 60, 1531–1543. (doi:10.1093/icb/icaa142)

8. Wellenreuther M, Svensson EI, Hansson B. 2014 Sexual selection and genetic colour polymorphisms in animals. Mol. Ecol. 23, 5398–5414. (doi:10.1111/mec.12935)

9. Carola V, Perlas E, Zonfrillo F, Soini HA, Novotny MV, Gross CT. 2014 Modulation of social behavior by the agouti pigmentation gene. Front Behav Neurosci 8, 259. (doi:10.3389/fnbeh.2014.00259)

10. Padilla SL et al. 2016 Agouti-related peptide neural circuits mediate adaptive behaviors in the starved state. Nat. Neurosci. 19, 734–741. (doi:10.1038/nn.4274)

11. Dijkstra PD, Maguire SM, Harris RM, Rodriguez AA, DeAngelis RS, Flores SA, Hofmann HA. 2017 The melanocortin system regulates body pigmentation and social behaviour in a colour polymorphic cichlid fish†. Proc. R. Soc. B: Biol. Sci. 284, 20162838. (doi:10.1098/rspb.2016.2838)

12. Jordan F, Babbitt KJ, Mclvor CC. 1998 Seasonal variation in habitat use by marsh fishes. Ecol. Freshw. Fish 7, 159–166. (doi:10.1111/j.1600-0633.1998.tb00182.x)

13. Maan ME, Sefc KM. 2013 Colour variation in cichlid fish: developmental mechanisms, selective pressures and evolutionary consequences. Semin Cell Dev Biol 24, 516–28. (doi:10.1016/j.semcdb.2013.05.003)

14. Cummings ME, Endler JA, Fuller H editor: RC. 2018 25 Years of sensory drive: the evidence and its watery bias. Curr. ZooDl. 64, 471–484. (doi:10.1093/cz/zoy043)

15. Seehausen O et al. 2008 Speciation through sensory drive in cichlid fish. Nature 455, 620–626. (doi:10.1038/nature07285)

16. Seehausen O, Alphen JJM van, Witte F. 1997 Cichlid Fish Diversity Threatened by Eutrophication That Curbs Sexual Selection. Science 277, 1808–1811. (doi:10.1126/science.277.5333.1808)

17. Kratochwil CF, Liang Y, Gerwin J, Woltering JM, Urban S, Henning F, Machado-Schiaffino G, Hulsey CD, Meyer A. 2018 Agouti-related peptide 2 facilitates convergent evolution of stripe patterns across cichlid fish radiations. Sci New York N Y 362, 457–460. (doi:10.1126/science.aao6809)

18. NüssleinLJVolhard C, Singh AP. 2017 How fish color their skin: A paradigm for development and evolution of adult patterns. Bioessays 39, 1600231. (doi:10.1002/bies.201600231)

19. Hoekstra HE, Robinson GE. 2022 Behavioral genetics and genomics: Mendel’s peas, mice, and bees. Proc. Natl. Acad. Sci. 119, e2122154119. (doi:10.1073/pnas.2122154119)

20. Maruska KP, Fernald RD. 2018 Astatotilapia burtoni: A model system for analyzing the neurobiology of behavior. Acs Chem Neurosci 9, 1951–1962. (doi:10.1021/acschemneuro.7b00496)

21. Korzan WJ, Robison RR, Zhao S, Fernald RD. 2008 Color change as a potential behavioral strategy. Horm Behav 54, 463–70. (doi:10.1016/j.yhbeh.2008.05.006)

22. Border SE, Piefke TJ, Fialkowski RJ, Tryc MR, Funnell TR, DeOliveira GM, Dijkstra PD. 2018 Color change and pigmentation in a color polymorphic cichlid fish. Hydrobiologia 832, 175–191. (doi:10.1007/s10750-018-3755-0)

23. Korzan WJ, Fernald RD. 2006 Territorial male color predicts agonistic behavior of conspecifics in a color polymorphic species. Behav Ecol 18, 318–323. (doi:10.1093/beheco/arl093)

24. Theis A, Roth O, Cortesi F, Ronco F, Salzburger W, Egger B. 2017 Variation of anal fin egg-spots along an environmental gradient in a haplochromine cichlid fish: BRIEF COMMUNICATION. Evolution 71, 766–777. (doi:10.1111/evo.13166)

25. Fernald RD, Hirata NR. 1979 The Ontogeny of Social Behavior and Body Coloration in the African Cichlid Fish Haplochromis burtoni. Zeitschrift Für Tierpsychologie 50, 180–187. (doi:10.1111/j.1439-0310.1979.tb01025.x)

26. Muske LE, Fernald RD. 1987 Control of a teleost social signal: I. Neural basis for differential expression of a color pattern. J Comp Physiology 160, 89–97. (doi:10.1007/bf00613444)

27. Muske LE, Fernald RD. 1987 Control of a teleost social signal: II. Anatomical and physiological specializations of chromatophores. J Comp Physiology 160, 99–107. (doi:10.1007/bf00613445)

28. Maruska KP, Fernald RD. 2010 Behavioral and physiological plasticity: rapid changes during social ascent in an African cichlid fish. Horm Behav 58, 230–40. (doi:10.1016/j.yhbeh.2010.03.011)

29. Maruska KP, Becker L, Neboori A, Fernald RD. 2013 Social descent with territory loss causes rapid behavioral, endocrine and transcriptional changes in the brain. J Exp Biology 216, 3656–66. (doi:10.1242/jeb.088617)

30. Whitaker KW, Alvarez M, Preuss T, Cummings ME, Hofmann HA. 2021 Courting danger: socially dominant fish adjust their escape behavior and compensate for increased conspicuousness to avian predators. Hydrobiologia, 1–15. (doi:10.1007/s10750-020-04475-9)

31. Egger B, Klaefiger Y, Theis A, Salzburger W. 2011 A Sensory Bias Has Triggered the Evolution of Egg-Spots in Cichlid Fishes. Plos One 6, e25601. (doi:10.1371/journal.pone.0025601)

32. Santos ME, Baldo L, Gu L, Boileau N, Musilova Z, Salzburger W. 2016 Comparative transcriptomics of anal fin pigmentation patterns in cichlid fishes. Bmc Genomics 17, 712. (doi:10.1186/s12864-016-3046-y)

33. Fernald RD, Hirata NR. 1977 Field study of Haplochromis burtoni: Quantitative behavioural observations. Anim Behav 25, 964–975. (doi:10.1016/0003-3472(77)90048-3)

34. Fernald RD. 1977 Quantitative behavioural observations of Haplochromis burtoni under semi-natural conditions. Anim Behav 25, 643–653. (doi:10.1016/0003-3472(77)90115-4)

35. Fang W, Blakkan D, Lee G, Bashier R, Fernald RD, Alvarado SG. 2022 DNA methylation of the endothelin receptor B makes blue fish yellow. bioRxiv, 2022.09.27.509821. (doi:10.1101/2022.09.27.509821)

36. Friard O, Gamba M. 2016 BORIS: a free, versatile openLJsource eventLJlogging software for video/audio coding and live observations. Methods Ecol Evol 7, 1325–1330. (doi:10.1111/2041-210x.12584)

37. Belleghem SMV, Papa R, OrtizLJZuazaga H, Hendrickx F, Jiggins CD, McMillan WO, Counterman BA. 2017 patternize: An R package for quantifying colour pattern variation. Methods Ecol Evol 9, 390–398. (doi:10.1111/2041-210x.12853)

38. Schindelin J et al. 2012 Fiji: an open-source platform for biological-image analysis. Nat Methods 9, 676–682. (doi:10.1038/nmeth.2019)

39. Weller HI, Belleghem SMV, Hiller AE, Lord NP. 2022 Flexible color segmentation of biological images with the R package recolorize. Biorxiv, 2022.04.03.486906. (doi:10.1101/2022.04.03.486906)

40. Gandrud C. 2013 Reproducible Research with R and R Studio. (doi:10.1201/b15100)

41. Cox S, Chandler S, Barron C, Work K. 2009 Benthic fish exhibit more plastic crypsis than non-benthic species in a freshwater spring. J. Ethol. 27, 497–505. (doi:10.1007/s10164-008-0148-2)

42. Hotta T, Kawasaka K, Satoh S, Kohda M. 2019 Fish focus primarily on the faces of other fish. Sci. Rep. 9, 8377. (doi:10.1038/s41598-019-44715-0)

43. Alcazar RM, Hilliard AT, Becker L, Bernaba M, Fernald RD. 2014 Brains over brawn: experience overcomes a size disadvantage in fish social hierarchies. J. Exp. Biol. 217, 1462–1468. (doi:10.1242/jeb.097527)

44. Howard MR, Ramsaroop MG, Hoadley AP, Jackson LR, Lopez MS, Saenz LA, Alward B. 2023 Female cichlids attack and avoid—but will still mate with—androgen receptor mutant males that lack male-typical body coloration. bioRxiv, 2023.11.02.565323. (doi:10.1101/2023.11.02.565323)

45. Li C-Y, Steighner JR, Sweatt G, Thiele TR, Juntti SA. 2021 Manipulation of the Tyrosinase gene permits improved CRISPR/Cas editing and neural imaging in cichlid fish. Sci. Rep. 11, 15138. (doi:10.1038/s41598-021-94577-8)

46. Liang Y, Meyer A, Kratochwil CF. 2020 Neural innervation as a potential trigger of morphological color change and sexual dimorphism in cichlid fish. Sci Rep-uk 10, 12329. (doi:10.1038/s41598-020-69239-w)

47. Dijkstra PD, Funnell TR, Fialkowski RJ, Piefke TJ, Border SE, Aufdemberge PM, Hartman HA. 2024 Sexual selection may support phenotypic plasticity in male coloration of an African cichlid fish. Proc. B 291, 20241127. (doi:10.1098/rspb.2024.1127)

48. Butler JM, Anselmo CM, Maruska KP. 2021 Female reproductive state is associated with changes in distinct arginine vasotocin cell types in the preoptic area of Astatotilapia burtoni. J Comp Neurol 529, 987–1003. (doi:10.1002/cne.24995)

49. Kohda M, Jordan LA, Hotta T, Kosaka N, Karino K, Tanaka H, Taniyama M, Takeyama T. 2015 Facial Recognition in a Group-Living Cichlid Fish. PLoS ONE 10, e0142552. (doi:10.1371/journal.pone.0142552)

