## Supplemental Table 1. Description of all behaviours scored with mean frequency and SEM. for "Facial and body colouration is linked to social rank in the African cichlid *Astatotilapia burtoni*"

| Behavior | Category | Description | Frequency Mean ± SEM |
| --- | --- | --- | --- |
| “Bite Male” | Agonistic | Snapping motion of the jaw of one male that touches another male. | 13.8 ±1.37 |
| “Bite female” | Agonistic/  Reproductive | Snapping motion of the jaw of a male that touches a female. | 25.0 ± 1.72 |
| “Chases female” | Agonistic/  Reproductive | A male moves toward a female and follows the female as it swims away. | 36.3 ±2.77 |
| “Chases male” | Agonistic | A male moves toward another male and follows the male as it swims away. | 21.1 ±2.03 |
| “Circle” | Reproductive | A male and a female carousel each other, typically inside the territory(the terracotta pot). | 3.27 ± 2.17 |
| “Dig pot/substrate” | Nesting | Fish takes up the gravel in its mouth and expels it outside the territory. | 30.8 ± 8.34 |
| “Entry/exit pot” | Locomotive | Fish enters or exits the territory (the terracotta pot) | 23.4 ±3.08 |
| “Flee” | Locomotive | Fish A quickly swims away from fish B in response to fish B | 26.9 ±2.31 |
| “Head-on-head fight” | Agonistic | Fish A and Fish B directly engage each other, maintaining a fixed orientation while moving back and forth in close proximity, their bodies aligned in a mirrored fashion. | 6.44 ± 2.33 |
| “Lateral side display” | Agonistic | Fish A adopts a lateral orientation, presenting its side to Fish B, often accompanied by fin extension and quivering movements. | 2.35 ± 0.33 |
| “Lead” | Reproductive | A male fish approaches a female, then reverses direction and guides the female toward the territory. The female follows, with the possibility of entering the territory. | 3.5 ±0.764 |
| “Quiver female” | Reproductive | A male fish performs rapid, tremulous movements next to a female | 4.12 ± 0.839 |
| “Quiver male” | Agonistic | A male fish performs rapid, tremulous movements next to another male. | 5.57 ± 0.586 |
